# The impact of fungal developmental structures on mechanical properties of mycelial materials

**DOI:** 10.1101/2025.04.01.644731

**Authors:** Kelsey Gray, Harley Edwards, Alexander G. Doan, Walker Huso, JungHun Lee, Wanwei Pan, Nelanne Bolima, Isha Gautam, Tuo Wang, Ranjan Srivastava, Marc Zupan, Mark R. Marten, Steven Harris

**Affiliations:** University of Maryland, Baltimore County (UMBC), Department of Chemical and Biochemical Engineering, 1000 Hilltop Circle, Baltimore, Maryland 21250, USA; Iowa State University, Department of Plant Pathology, Entomology and Microbiology, Ames, IA 50011, USA; Michigan State University, Department of Chemistry, 578 S. Shaw Ln, East Lansing, MI 48824, USA; University of Connecticut, Department of Chemical and Biomolecular Engineering, Storrs, CT 06269, USA; University of Maryland, Baltimore County (UMBC), Department of Mechanical Engineering, 1000 Hilltop Circle, Baltimore, Maryland 21250, USA

## Abstract

This study explores how suppressing asexual development in *Aspergillus nidulans* enhances the mechanical properties of mycelial materials. Using four aconidial mutants (Δ*brlA*, Δ*flbA*, Δ*fluG*, and *fadA*^*G42R*^) that lack asexual development and a control strain (A28) that undergoes typical asexual development, we found that the absence of asexual development significantly improves mechanical strength. All mutants exhibited higher ultimate tensile strength (UTS) than the control, with Δ*fluG* and Δ*brlA* (fluffy nonsporulating, FNS phenotype) showing the highest UTS. Additionally, *fadA*^*G42R*^ and Δ*flbA* (fluffy autolytic dominant, FAD phenotype) demonstrated significantly higher strain at failure (SF), linked to increased autolysis and lower dry cell mass compared to the control and FNS mutants. Solid-state NMR analysis revealed that autolysis in FAD mutants disrupts galactofuranose-related metabolic processes, altering cell wall composition and contributing to higher elasticity. These findings suggest that suppressing asexual development enhances mycelial material strength, while autolysis mechanisms influence elasticity. This research highlights the potential for genetic manipulation in fungi to engineer advanced mycelial-based materials with tailored mechanical properties.

## 1. INTRODUCTION

The urgent need for sustainable materials continues to grow, as environmental concerns escalate and non-renewable resources become increasingly scarce ^[1]^. Among emerging alternatives to traditional, non-biodegradable materials, fungal-based biomaterials stand out for their unique combination of advantages: they are fully bio-based, cost-effective to manufacture, and naturally biodegradable ^[2]^. The versatility of fungi enables the production of materials with diverse properties, leading to successful applications across multiple industries, from textiles and packaging to automotive components and building materials ^[3]^. At the heart of these innovations is the mycelium, the vegetative structure of fungi, which forms intricate networks of microscopic filaments characterized by remarkably efficient growth patterns and robust cell wall architecture ^[4]^. These biological features translate into impressive mechanical capabilities - depending on the fungal species and growth substrate selected, mycelium-based materials can match or exceed the mechanical properties of conventional materials ^[5]^.

Recent research has revealed that the mechanical properties of mycelial materials are intricately linked to specific fungal characteristics, including hyphal diameter, cell wall composition, and hyphal packing density [3]. This understanding opens up exciting possibilities for precise material engineering through genetic manipulation of fungi, potentially allowing scientists to fine-tune these characteristics for specific applications. However, to fully realize this potential, a deeper understanding is needed regarding the complex relationships between fungal genetics, resulting hyphal composition, morphology, and mechanical properties. Such research would enable more precise control over material properties and accelerate the development of next-generation sustainable materials.

In our previous work, we explored the impact on mycelial-material mechanical properties of deleting the gene encoding the last protein-kinase (*mpkA*) in the cell wall integrity (CWI) signaling pathway in *Aspergillus nidulans* ^[6]^. The CWI pathway is responsible for cell wall repair, and deletion of *mpkA* leads to significantly altered mycelial phenotypes ^[7]^. We hypothesized that the Δ*mpkA* mutant would have cell walls with altered composition, that these would be weaker, and that this would lead to significantly weaker mycelial materials. In contrast, we found the Δ*mpkA* mutant generated significantly (approximately 55%) stronger (ultimate tensile strength, UTS), and more elastic (strain at failure, SF), mycelial material. To determine the cause of this behavior, we carried out an extensive phenotype analysis of the Δ*mpkA* mutant. While the composition of the walls is somewhat different, a more striking difference appears to be the lack of developmental structures. This appears to result in more densely packed hyphae and stronger resultant material. This finding was consistent with work from others ^[8]^.

These observations led us to hypothesize that fungal strains devoid of asexual developmental features would, in-general, lead to stronger mycelial materials. Asexual development in filamentous fungi is a complex process, regulated by a host of different gene products ^[9]^. When filamentous fungi grow vegetatively, hyphae typically extend at the tip, which is where nutrient assimilation primarily occurs ^[10]^. In response to nutrient deprivation and other signals, these fungi undergo asexual development to produce conidia ^[11]^, which serve as protective structures, ready to germinate and grow when additional nutrients become available ^[12]^. Thus, vegetative growth occurs prior to the onset of asexual development ^[11, 13, 14]^, which creates morphological changes ^[15]^, impacting both hyphal biomass and the density of hyphae in a given area ^[10, 16]^. These factors appear to play a role in determining the mechanical properties of mycelial materials ^[17]^.

To test the hypothesis that the absence of asexual development leads to stronger mycelial materials, we used the model fungus *Aspergillus nidulans*. We chose four different *A. nidulans* mutants (Δ*brlA*, Δ*flbA*, Δ*fluG, and fadA*^*G42R*^) that have previously been shown to not undergo asexual development (i.e., aconidial mutants ^[18-20]^) and a wild type (A28) control which undergoes normal asexual development ^[21, 22]^. Within the group of aconidial, or “fluffy” mutants (based on their appearance when grown on a solid surface), two phenotypes are represented ^[23]^. The “fluffy non sporulating” (FNS) phenotype, (i.e., Δ*fluG*, Δ*brlA*) exhibit exclusively vegetative growth, but proliferate at different rates, while the “fluffy autolytic dominant” (FAD) mutants (Δ*flbA, fadA*^*G42R*^) resemble FNS mutants, with the addition of a pronounced and accelerated autolysis phenotype ^[23]^. Our testing produced data which supports our hypothesis, as we find suppression of asexual development leads to mycelial material with increased ultimate tensile strength (UTS).

## 2. MATERIALS & METHODS

### 2.1 Fungal Strains and Growth Media

The *Aspergillus nidulans* strains used in this study were all obtained from the Fungal Genetic Stock Center (FGSC, Manhattan KS, USA) ^[22]^ and are listed in Table 1. Initially, strains were grown on modified MAG-V agar plates, which consisted of 2 g/L BD Bacto Peptone, 1 mL/L vitamin solution, 1 mL/L Hutner’s trace elements solution, 20 g/L granulated agar, 1.12 g/L uracil, 1.22 g/L uridine, 35.065 g/L NaCl, 20 g/L glucose, and 20 g/L malt extract ^[6, 24]^. Hutner’s trace element solution was prepared with the following components: 22 g/L ZnSO_4_·7H_2_O, 11 g/L H_3_BO_3_, 5 g/L MnCl_2_·4H_2_O, 5 g/L FeSO_4_·7H_2_O, 1.7 g/L CoCl_2_·6H_2_O, 1.6 g/L CuSO_4_·5H_2_O, 1.5 g/L Na_2_MoO_4_·2H_2_O, and 50 g/L EDTA (Na4). The vitamin solution included 100 mg/L each of biotin, pyridoxine, thiamine, riboflavin, p-aminobenzoic acid, and nicotinic acid ^[25]^. For cultivating mycelial materials, modified YGV medium was utilized. This medium, per liter, contained 1 g/L yeast extract, 2 g/L BD Bacto peptone, 1 g/L Bacto casamino acids, 1 mL/L vitamin solution, 1 mL/L Hutner’s trace elements solution, 44.21 g/L KCl, 1.12 g/L uracil, 1.22 g/L uridine, 20 g/L glucose, 20 g/L malt extract, 10 g/L proline, 50 mL/L nitrate salts, and 5 mL/L MgSO4 solution ^[6, 24]^. The nitrate salts were composed of 142.7 g/L KNO_3_, 10.4 g/L KCl, 16.3 g/L KH_2_PO_4_, and 20.9 g/L K_2_HPO_4_, while the MgSO4 solution required 104 g/L of MgSO4.

**Table 1.**
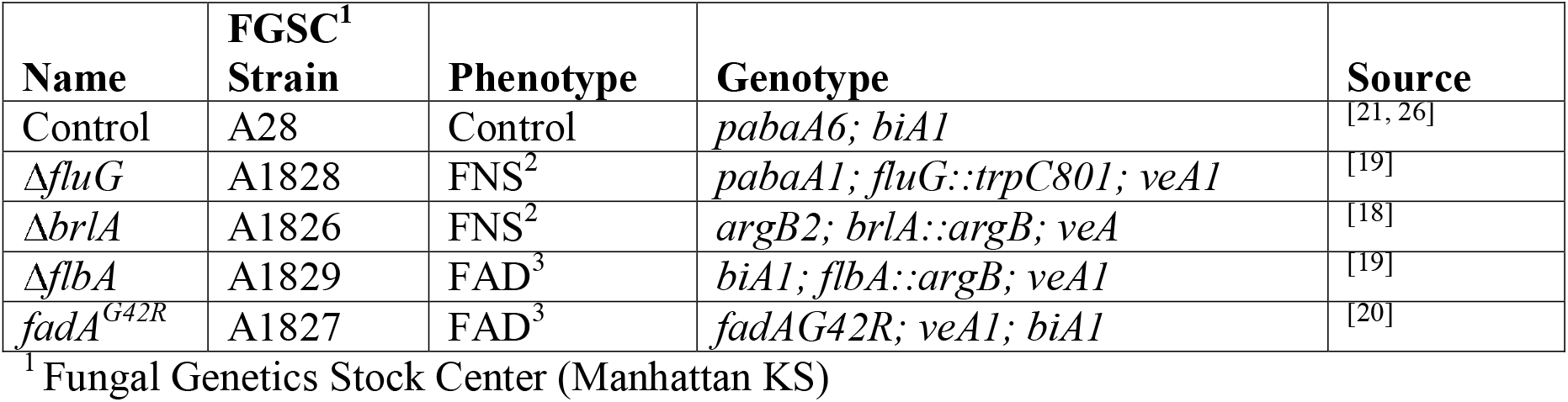
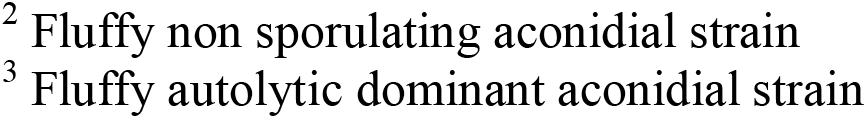
Aspergillus nidulans strains used in this study.

| Name | FGSC<sup>1</sup><br>Strain | Phenotype | Genotype | Source |
| --- | --- | --- | --- | --- |
| Control | A28 | Control | *pabaA6; biA1* | [21, 26] |
| $\Delta fluG$ | A1828 | FNS<sup>2</sup> | *pabaA1; fluG::trpC801; veA1* | [19] |
| $\Delta brlA$ | A1826 | FNS<sup>2</sup> | *argB2; brlA::argB; veA* | [18] |
| $\Delta flbA$ | A1829 | FAD<sup>3</sup> | *biA1; flbA::argB; veA1* | [19] |
| $fadA^{G42R}$ | A1827 | FAD<sup>3</sup> | *fadAG42R; veA1; biA1* | [20] |

<sup>1</sup> Fungal Genetics Stock Center (Manhattan KS)
<sup>2</sup> Fluffy non sporulating aconidial strain
<sup>3</sup> Fluffy autolytic dominant aconidial strain

### 2.2 Growing Mycelial Materials

The procedure for generating mycelial material from the control strain has been described previously ^[6]^. Briefly, fungi were cultivated on modified MAG-V plates to produce conidia, which were then collected, quantified, and used to inoculate YGV medium in a 60 mm petri dish (Fisherbrand). The dish was incubated at 28□ for 120 hours. After incubation, the grown material disc was harvested and rinsed sequentially with bleach and deionized (DI) water. To prepare material from aconidial mutants, the process began by extracting two “agar cores” from fungi grown on modified MAG-V plates. These cores were placed into a cryogenic vial containing 0.5 mL of 3 mm diameter glass beads, followed by the addition of 0.9 mL of DI water. The vial was homogenized for 10 seconds at 5000 rpm using a Mini Beadbeater (Biospec Products). A 200 µL aliquot of the homogenized solution was then transferred to a 60 mm petri dish (Fisherbrand) containing 7.5 mL of modified YGV medium and incubated at 28□ for 120 hours. The resulting material disc was harvested and washed with bleach and DI water.

### 2.3 Scanning Electron Microscopy (SEM)

Scanning electron microscopy (SEM) was performed following the methodology outlined in ^[6]^. In summary, mycelial samples were examined both prior to, and following, tensile testing. The samples were mounted on an aluminum stub with double-sided carbon tape, coated with a thin layer of gold/palladium using a Cressington 108 Manual Sputter Coater, and then analyzed using an FEI Nova NanoSEM 450.

### 2.4 Asexual Development (Conidiation)

For quantification of conidia, mycelial-material was grown as above, and soaked in hexamethyldisilazane (HMDS) for 5 min, then air dried for at least 90 minutes ^[6]^. The dried material was added to a cryogenic vial containing 3 glass beads (diameter 3 mm), and homogenized with a Mini Beadbeater (Biospec Products) at 5000 rpm for 5 seconds. Then 1 mL of DI water was added, vortexed for 15 seconds, homogenized again at 5000 rpm for 5 seconds, then vortexed once more for 15 seconds. Homogenized samples were diluted with DI water before being vortexed and placed onto a hemocytometer for counting conidia.

### 2.5 Tensile Testing (Mechanical Testing)

Tensile testing was performed on 4 × 40 mm coupons, which were laser-cut (Universal Laser Systems VLS 3.6/6) from mycelial disks. A minimum of five coupons were prepared from each disk, with three disks tested per genotype. Prior to testing, the coupons were immersed in a 40:60 glycerol-to-water solution for 1 minute ^[27]^, coated in petroleum jelly ^[28]^, and dried using warm air for about 1 minute per side. The dimensions of the coupons were accurately determined using a dissection microscope (Leica Microscope s9i Model meb115), and they were attached to a paper frame for stability using cyanoacrylate adhesive (Loctite) and NaHCO_3_ powder (to speed up drying ^[29]^). The frame was secured to the tensile testing machine (Instron 3369) with double-sided tape. Just before testing, the paper frame was cut on both sides to ensure only the mycelial material was subjected to tensile stress. Reflective tape was applied to the paper frame to facilitate strain measurement via a laser extensometer (Electronic Instrument Research Model LE-01) during the application of stress. Stress was recorded using a sensor (Transducer Techniques Sensor) connected to the tensile testing device. Testing was manually halted once material failure occurred ^[6]^.

### 2.6 Cell Wall Composition

The proportion of rigid and mobile components was assessed by examining peak volumes in 2D ^13^C-^13^C spectra, utilizing 53 ms CORD and DP refocused J-INADEQUATE techniques ^[30]^, respectively. Solid-state NMR data were acquired using an 800 MHz Bruker Avance Neo NMR spectrometer equipped with a 3.2 mm HCN magic-angle spinning probe. The ^13^C chemical shifts were referenced to the tetramethylsilane (TMS) scale. Peak volumes were quantified using the integration tool in Bruker Topspin 4.1.4 software. To reduce errors caused by overlapping signals, only distinct and well-separated peaks were included in the analysis following a protocol recently established for fungal cell wall compositional analysis ^[31]^. The assigned NMR peaks and corresponding integral volumes were represented as a bar graph using Microsoft Excel.

### 2.7 Significance Testing

All data were subjected to T-tests to identify any significant differences between groups using Microsoft Excel. Cell wall composition measurements were normalized to dry cell mass (DCM) for each strain to account for differences in biomass. The normalized data were analyzed using IBM SPSS Statistics (version 30). For each cell wall component (rigid phase: β-glucan, chitin, α -glucan; mobile phase: galactofuranose, galactose, β-glucan, α-mannose 1,2, α-mannose O, α-mannose 1,6), a one-way analysis of variance (ANOVA) was performed with strain type as the independent variable and the normalized measurement (Measurement/DCM) as the dependent variable. Homogeneity of variance was assessed using Levene’s test. Tukey’s Honestly Significant Difference (HSD) post-hoc test was employed to analyze pairwise differences between fungal strains for each cell wall component. Statistical significance was set at *P* < 0.05. This approach allowed for identification of strain-specific differences in cell wall composition across both rigid and mobile phases, providing insight into how genetic variation between FAD and FNS mutants influence cell wall architecture.

## 3. RESULTS AND DISCUSSION

### 3.1 Fungal Morphology

To confirm the presence or absence of asexual developmental structures (i.e., conidia), mycelial material (*n* = 2 coupons per genotype) was imaged before tensile testing, using scanning electron microscopy (SEM). **Figure 1A** shows there was an abundance of conidia in material generated from the control strain (A28), but this is not seen in the material generated by any of the four aconidial mutants (**Figures 1B-E)**.

**Figure 1.**
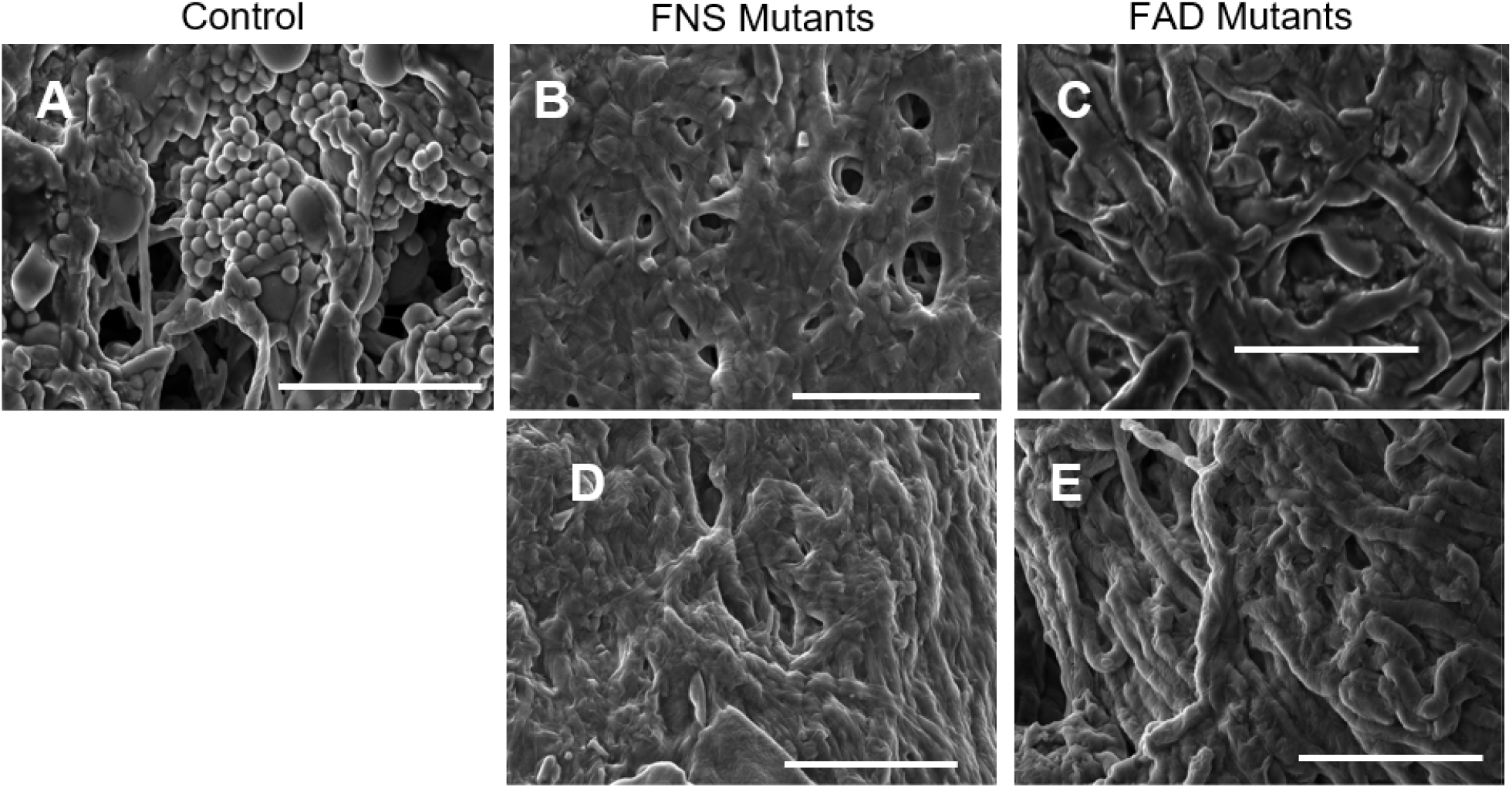
Representative SEM images of mycelial material show three-dimensional structure and morphological features. (**A**) Control (A28) sample surface showing an abundance of developmental structures (e.g., conidia), loose packing of hyphae, and voids in the material. Aconidial mutants: (B) Δ*fluG*, (C) Δ*brlA*, (D) Δ*flbA* and (E) *fadA*^G42R^, surfaces show more densely packed hyphae, fewer voids, and no developmental structures. All bars = 30 µm.

These images confirm findings from previous studies ^[32]^ regarding the degree of asexual development in each of the strains represented. We note that in regard to developmental structures, SEM images of freshly harvested mycelial material (**Figure S1**) was similar in appearance to that found in the material used for tensile testing. After tensile testing, SEM images of the fracture point for each genotype were collected (**Figure 2**) and show similar phenomena. The control strain has abundant conidia, while all the other strains do not.

**Figure 2.**
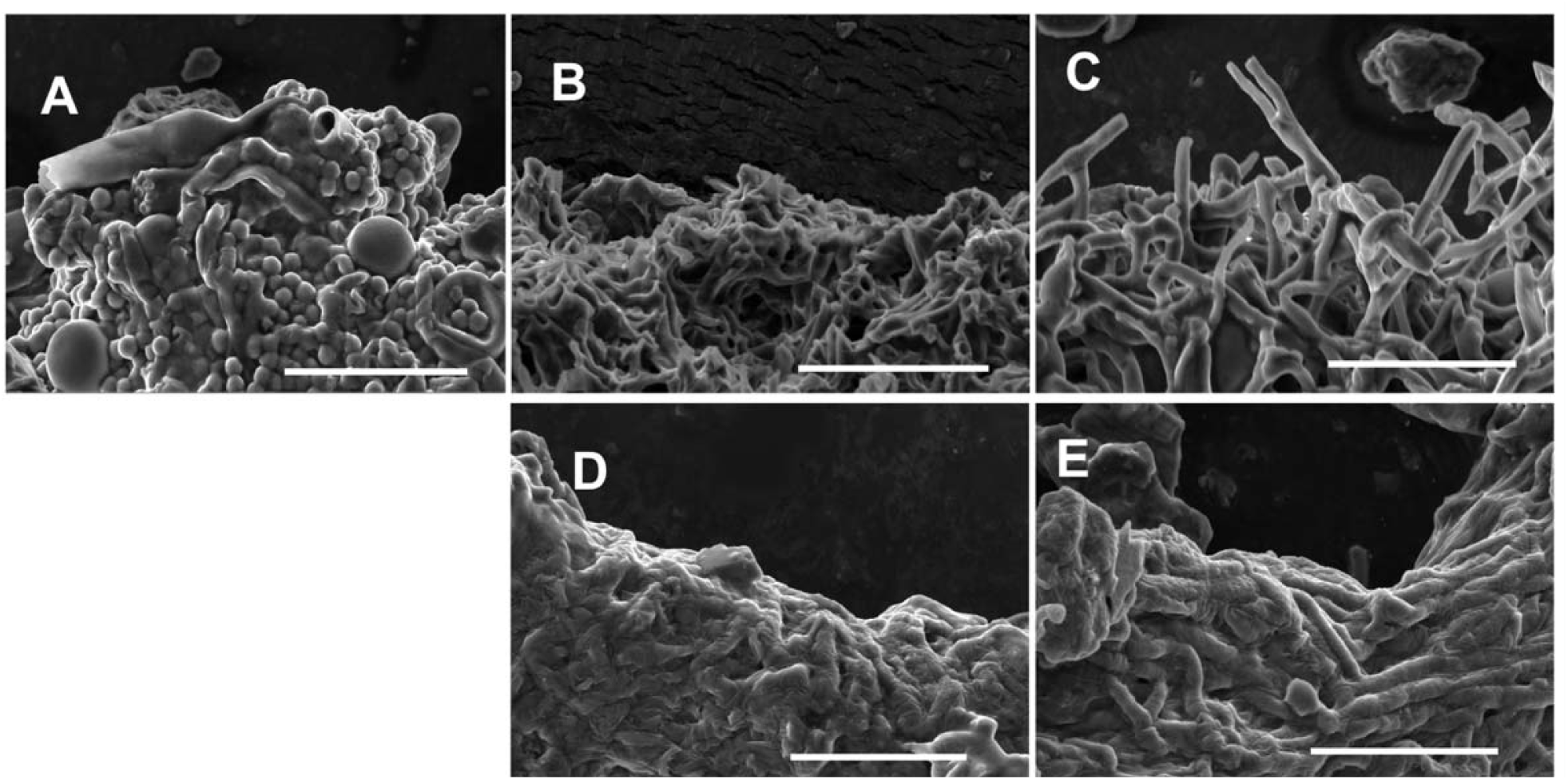
Representative SEM images of mycelial material three-dimensional structure and morphological features at the fracture point after tensile testing. (**A**) Control (A28) sample shows an abundance of developmental structures (e.g., conidia) present. Aconidial mutants: (B) Δ*fluG*, (C) Δ*brlA*, (D) Δ*flbA* and (E) *fadA*^G42R^ show exclusively hyphal morphology with no developmental structures. All bars = 30 µm.

Further evidence of the difference in asexual development between the control and mutant strains is provided in **Figure 3**, where the control (*n* = 2 per genotype) produced on average 2.31E10 conidia/gram, compared to all of the mutants which produced approximately 10^4^ fewer conidia/g (*P < 0*.*05*).

**Figure 3.**
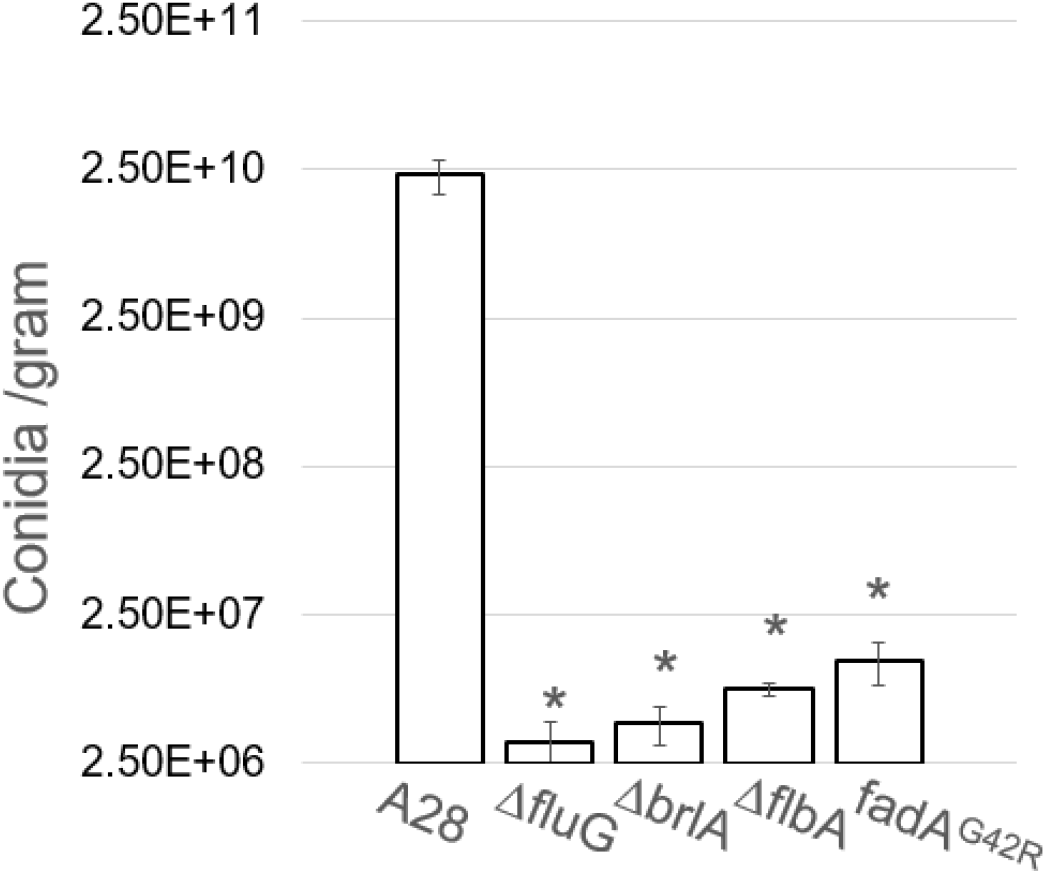
Average number of conidia produced per gram of mycelial material for each genotype tested. Star (*) indicates significant difference (*P* ≤ 0.05) when compared to control (A28).

### 3.2 Mechanical Properties

To determine material mechanical properties, a mycelial-material disc was prepared as described above and cut into five strips. Each of these strips was tensile tested, and a stress-strain curve from a typical test is shown in **Figure 4**. The numbers in the figure indicate relative position of the tested coupons cut from the mycelial material disc. Positions 1-5 were immediately adjacent to each other, with position 3 in the middle of the material disk. Similar to results in **Figure 4**, all tensile testing showed only random variation with regard to position, implying material was homogeneous with regard to material properties. For each genotype, three mycelial-material discs were tested (*n* = 15 strips per genotype). In all tests, materials displayed linear elasticity prior to failure, a trait indicative of brittle materials ^[33]^.

**Figure 4.**
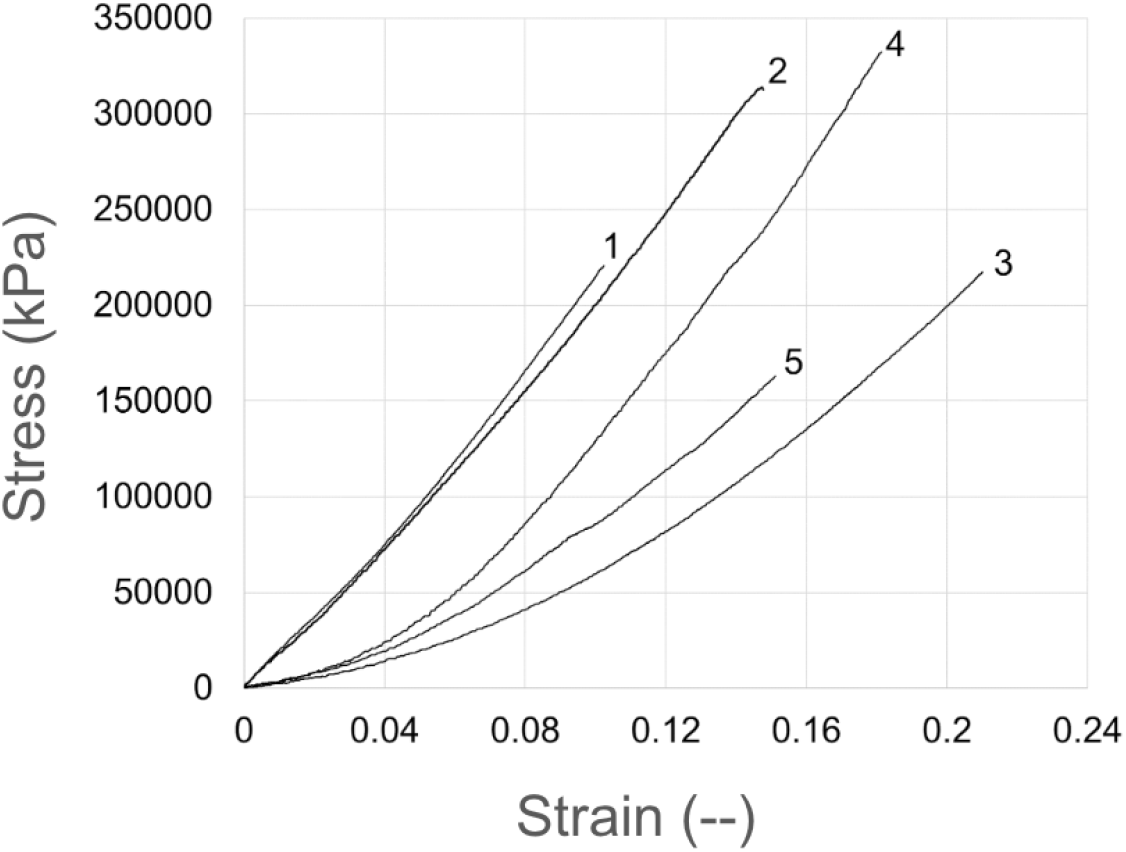
Typical stress-strain curves for mycelial material generated from the control (A28) strain. Five test strips were cut from the center of a mycelial material disk, numbers on the graph indicate relative position of strips. This material shows linear elasticity before failure.

Data collected from the tensile testing was used to determine: ultimate tensile strength (UTS), and strain at failure (SF). Average values for each of these properties are shown in **Figure 5. Figure 5A** shows that compared to the control (UTS of 158 ± 24 kPa), all mutants, Δ*fluG*, Δ*brlA*, Δ*flbA*, and *fadA*^*G42R*^, have significantly higher (*P* ≤ 0.05) UTS of 323 ± 33 kPa, 306 ± 8 kPa, 218 ± 17 kPa, and 288 ± 30 kPa respectively. Our data show the aconidial mutants are mechanically stronger than the control, which suggests that asexual development weakens the mechanical strength of mycelial materials. **Figure 5B** shows the control, with a SF of 0.12 ± 0.01, wa statistically similar (*P* ≥ 0.1) to each FNS mutant, Δ*fluG* with an SF of 0.11 ± 0.01 and Δ*brlA* with an SF of 0.12 ± 0.01. In contrast, the FAD mutants, *fadA*^*G42R*^ and Δ*flbA*, showed significantly higher SF when compared to the control with a SF of 0.20 ± 0.02 and 0.21 ± 0.01 respectively (*P* ≤ 0.05).

**Figure 5.**
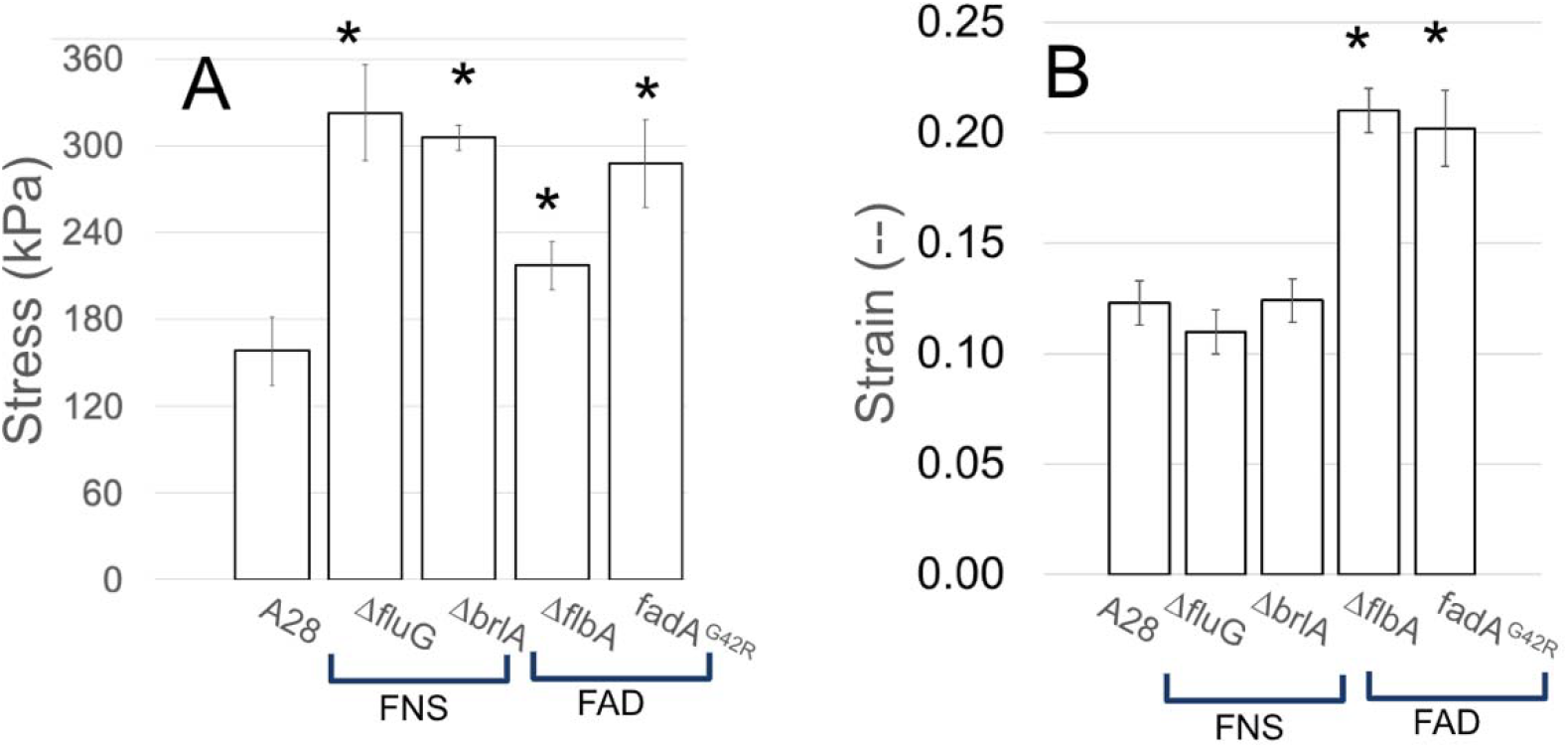
Mechanical properties of mycelial materials. Measurements made during tensile testing (*n* = 15 for each fungal strain) of material generated from control (A28) and aconidial mutants (Δ*brlA*, Δ*flbA*, Δ*fluG*, and *fadA*^G42R^). Averages of (**A**) ultimate tensile strength (stress kPa), and (B) strain at failure (−-). Star (*) indicates significant difference (*P* ≤ 0.05) when compared to control (A28).

Our work evaluates the mechanical properties of mycelial materials generated from aconidial fungal mutants, and finds that the inhibition of asexual development appears, in general, to increase the mechanical strength of mycelial materials. Our data suggest that the phenotypes of the aconidial mutants may have resulted in a hierarchy of strength among mycelial materials with FNS mutants producing the strongest materials, followed by the FAD mutants, and finally, the control. Mycelial materials generated from the FAD mutants were found to be the most elastic based on their increased strain at failure (**Figure 5B**). Considering that a major phenotype expressed exclusively among the FAD mutants are their significant degree of autolysis ^[23]^, we hypothesized that the accelerated autolysis occurring in these strains may have contributed to altered mechanical properties.

### 3.3 Confirmation of Autolysis and Impact on Cell Wall Composition

The degree of autolysis exhibited in all strains was assessed by comparing the dry cell mas (DCM) of the mycelial materials generated for each genotype. We hypothesized that strains with the autolytic phenotypes (i.e., FAD mutants) would show a reduced DCM, consistent with previous findings ^[34]^. The results in **Figure 6** show this is the case. The FAD mutants have significantly lower DCM when compared to either the control or the FNS mutants, despite the FAD mutants having a hyphal-dominant morphology which typically leads to increased amounts of biomass ^[13]^. The significantly reduced biomass in both FAD mutants suggests that accelerated autolysis may be the phenotypic mechanism for their increased strain at failure (SF) observed during mechanical testing.

**Figure 6.**
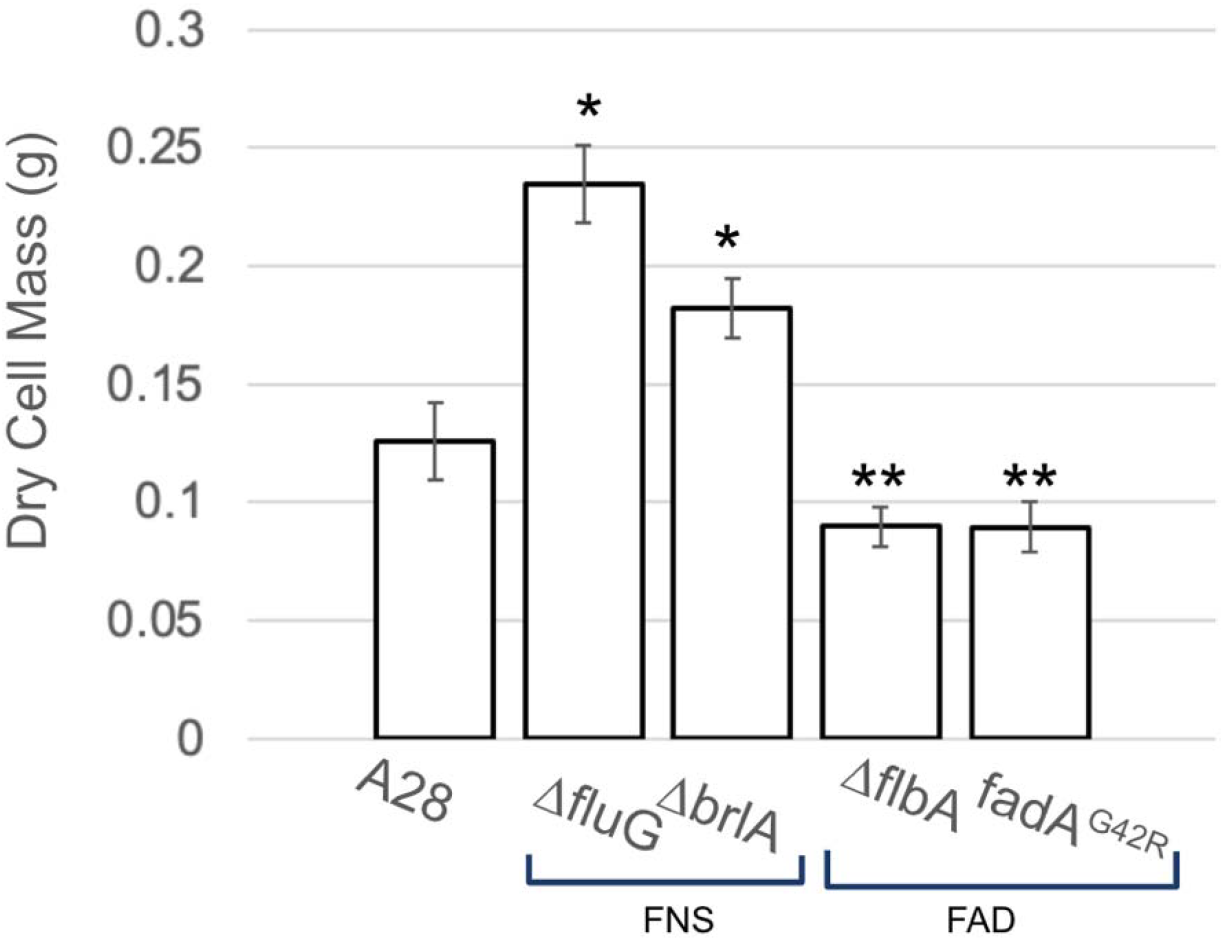
Average dry cell mass of mycelial materials generated from each genotype. Star (*) indicates significant difference (*P* ≤ 0.05) when compared to control (A28). Double star (**) indicates significant difference (*P* ≤ 0.05) whe compared to control and FNS mutants (Δ*fluG* and Δ*brlA*).

To determine if autolytic degradation in FAD mutants led to altered cell wall composition we used a high-resolution, solid-state NMR approach we have used previously ^[6]^ which assessed composition of both the rigid-core and mobile-domain of the cell wall. Measurements were normalized by DCM (**Figure 6**) and results are shown in **Figure 7**. While there is significant variation in the data, statistical analysis of normalized cell wall components provides insight and revealed distinct patterns of alteration across fungal strains. For example, in the rigid phase, the Δ*flbA* mutant exhibited significantly higher β-glucan content compared to all other strains (*P* < 0.001), with levels approximately 4-5 fold higher than the control strain. Neither chitin nor α-glucan showed statistically significant differences between strains, though Δ*flbA* displayed numerically higher values for both components.

**Figure 7.**
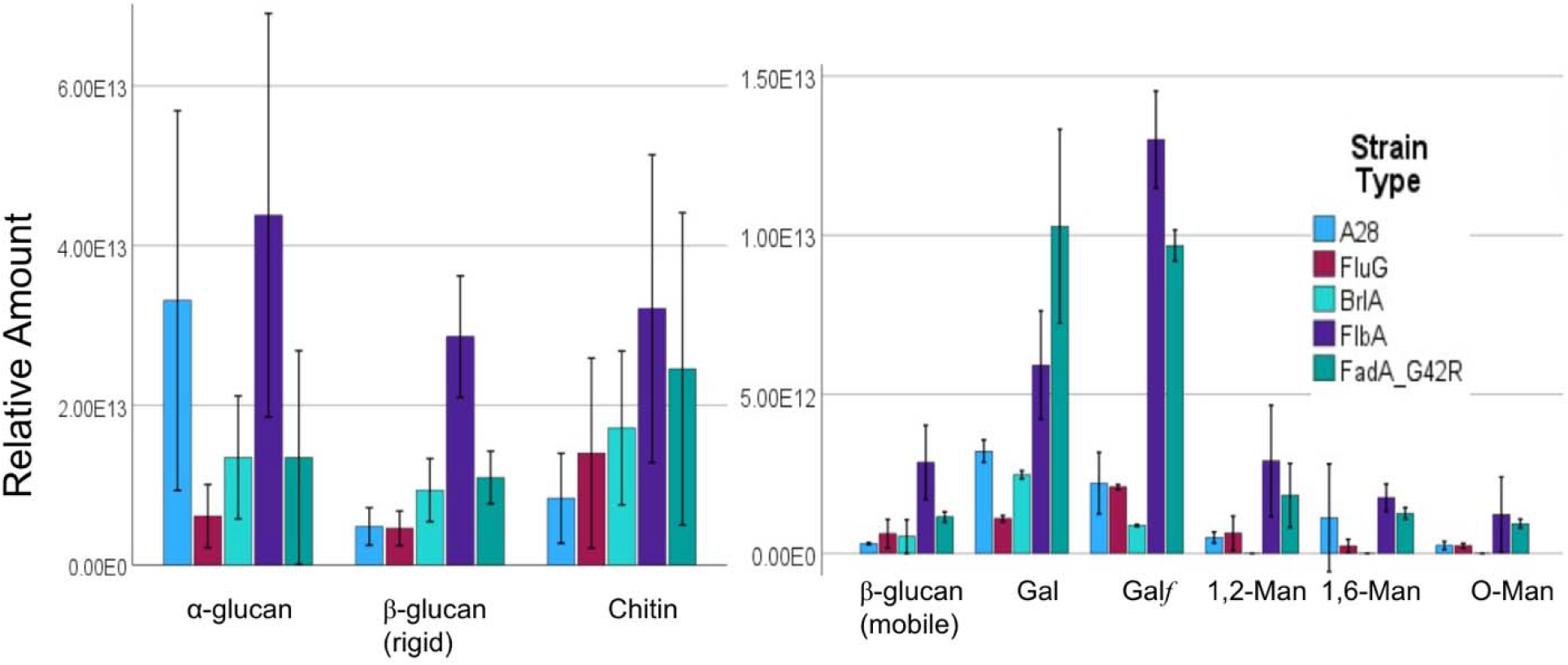
Results from ssNMR cell wall analysis. Left: Rigid phase, Right: Mobile phase. Significance testing described in the text.

In the mobile phase, component-specific effects were observed across mutant strains. For galactofuranose (Gal*f*), both FAD mutants showed significantly elevated levels compared to the control and FNS mutants, with Δ*flbA* exhibiting the highest levels (*P* < 0.001), followed by *fadA*^*G42R*^ (*P* < 0.001). The content of the mobile phase β-glucan was significantly higher in Δ*flbA* compared to A28, Δ*brlA*, and Δ*fluG* (*P* < 0.05). Interestingly, for galactose (Gal), *fadA*^*G42R*^ demonstrated the highest levels, significantly exceeding those in the control and FNS mutants (*P* < 0.01). The mannose derivatives (1,2-Man, O-Man, 1,6-Man) displayed limited significant differences between strains.

These results reveal genotype-specific, rather than phenotype-specific (i.e., FAD, FNS), effects on cell wall composition. The Δ*flbA* mutation consistently displayed increased glucan content in both rigid and mobile β-glucan phases, while also significantly elevating galactofuranose levels. *fadA*^*G42R*^, despite being categorized with Δ*flbA* as an FAD mutant, demonstrated a more selective effect, primarily increasing galactose-containing components (galactofuranose and galactose). In contrast, FNS mutants (Δ*fluG* and Δ*brlA*) showed minimal alterations in cell wall composition compared to the control strain.

These findings suggest that disruption of the Δ*flbA* gene fundamentally alters cell wall architecture through increased deposition of glucans and galactofuranose, likely contributing to substantial changes in mechanical properties. The differential effects observed between Δ*flbA* and *fadA*^*G42R*^ indicate these genes may influence cell wall composition through distinct mechanisms, despite belonging to the same regulatory pathway as represented through the figure created by Son et al ^[9]^. The component-specific nature of these alterations highlights the complex relationship between genetic variation and resulting phenotypic changes in fungal cell wall structure. These distinct compositional signatures may explain previously observed differences in mechanical properties between these mutant strains.

## 4. Conclusion

Consistent with our initial hypothesis, we found that mycelial materials grown from *A. nidulans* aconidial mutants exhibited statistically significant increases in ultimate tensile strength. Additionally, when compared to the control and FNS mutants, both FAD mutants displayed a statistically significant increase in the strain at failure. This appears to be due to their accelerated autolysis, altering metabolic processes - with an emphasis on processes involving galactofuranose, leading to altered cell wall composition.

## Acknowledgements

We gratefully acknowledge significant assistance from Dr. Tagide deCarvalho in the UMBC, Keith R. Porter Imaging Facility regarding SEM preparation and imaging; Drs. Michael Duffy and Joao Santos (Engineering Testing, LLC) for their assistance and guidance with material tensile testing; Dr. Govind Rao (UMBC, Center for Advanced Sensor Technology) for use of the laser cutter to prepare samples; Drs. Jordan Baumbach and Tagide deCarvalho for general guidance with this project.

## Supplemental Figures

**Figure S1.**
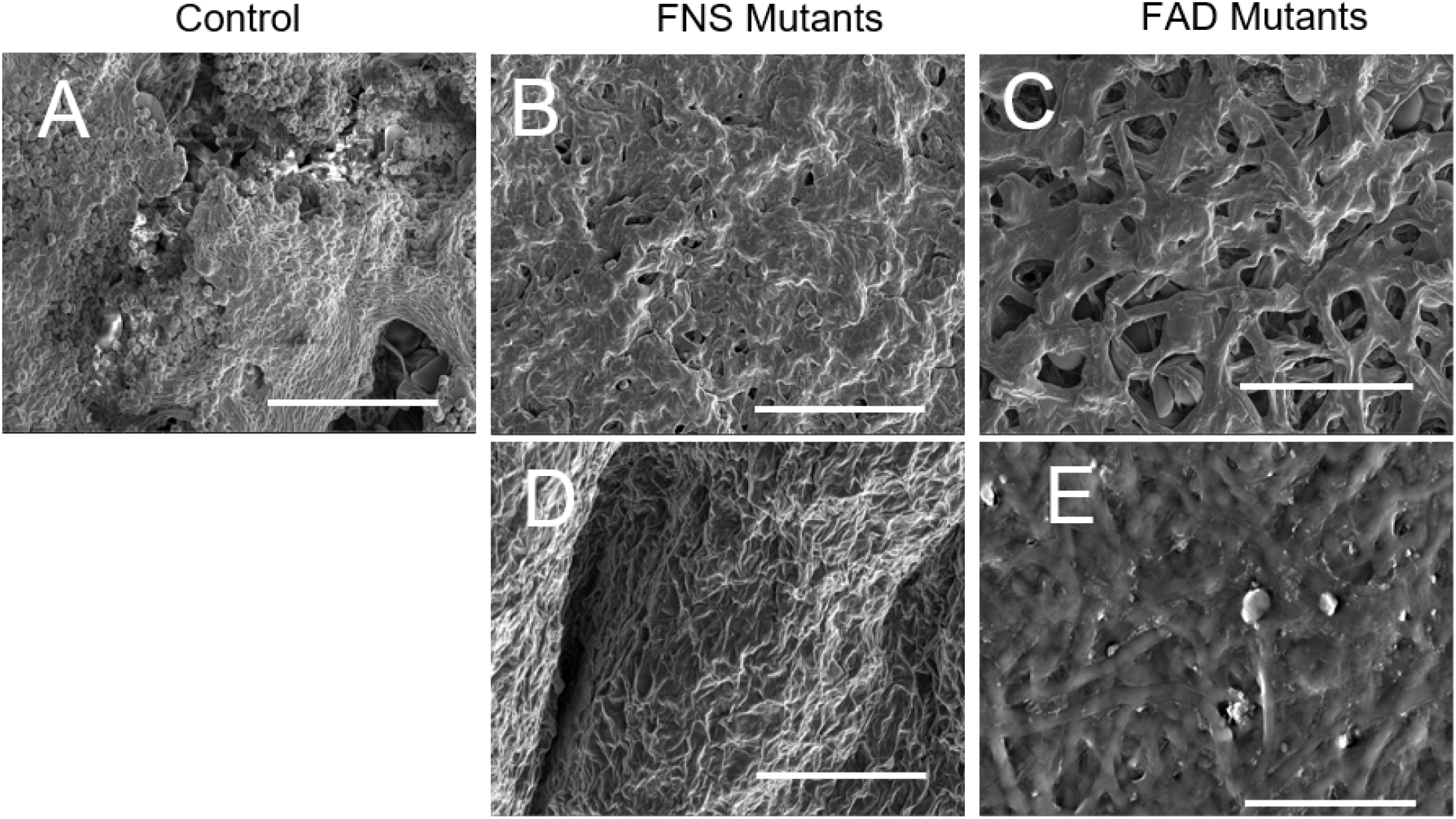
Representative SEM images of freshly harvested mycelial material three-dimensional structure and morphological features (some collapsing is present). (**A**) Control (A28) sample surface showing an abundance of asexual developmental structures (i.e., conidia), loose packing of hyphae, and voids in the material. Aconidial mutants: (B) Δ*fluG*, (C) Δ*brlA*, (D) Δ*flbA* and (E) *fadA*^G42R^, do not show any aconidial development structures (i.e., conidia). All bars = 50 µm.

